# Disrupted dynamics of brain structure–function coupling link genetic risk of Alzheimer’s Disease and aging to cognitive decline in 34,067 adults

**DOI:** 10.64898/2026.01.06.697897

**Authors:** Xinran Wu, Zhengxu Lian, Songjun Peng, Yu Liu, Nanyu Kuang, Gechang Yu, Zhaowen Liu, Jianfeng Feng, Jie Zhang

## Abstract

Understanding how a stable structural connectome supports flexible cognition, especially in aging, is a fundamental question in neuroscience. While static structure–function coupling (SFC) is well-studied, the dynamic decoupling of functional activity from structural constraints—dynamic SFC (DSFC)—remains poorly understood. Leveraging MRI data from 34,067 UK Biobank participants (ages 45–82), we characterized the distinct roles of SFC and DSFC in aging, cognition, and health. We found that SFC and DSFC followed spatially divergent aging trajectories: SFC declined primarily in sensorimotor systems, whereas DSFC decreased most prominently in higher-order networks. Both SFC and DSFC in higher-order networks were positively correlated with cognitive performance (e.g., fluid intelligence). However, the associations with mental and physical health diverged between the two measures: reduced DSFC was predominantly linked to health burdens in high-order default/limbic networks, whereas weakened SFC was primarily associated with health burdens in low-order sensory-motor networks. Genetic analyses revealed that Alzheimer’s risk, specifically APOE ε4 dosage, significantly reduced DSFC in higher-order cognitive networks and SFC in visual cortex. Mediation analyses further demonstrated that aging and APOE ε4-linked cognitive decline were mediated via visual SFC and ventral attention DSFC. These findings position static and dynamic coupling as complementary mechanisms: static SFC preserves network robustness, while dynamic SFC enables transient reconfiguration for complex integration. Together, our results highlight these dual mechanisms as crucial for maintaining cognitive flexibility, providing potential biomarkers for age-related neurodegeneration.

## Introduction

One of the central challenges in neuroscience is understanding how a relatively stable structural connectome gives rise to flexible and adaptive brain functional dynamics, a key feature that supports complex behaviors and higher cognitive functions (Honey, Thivierge, & Sporns, 2010; Ju & Bassett, 2020; Park & Friston, 2013; Uddin, 2013). This characteristic is captured by structure-function coupling (SFC) (Fotiadis et al., 2024; Suárez, Markello, Betzel, & Misic, 2020), which quantifies the constraints that structural connection (SC) impose on the synchrony of functional activity, or functional connection (FC) at the regional level (Fotiadis et al., 2023; Gu, Jamison, Sabuncu, & Kuceyeski, 2021; Preti & Van De Ville, 2019). The brain’s ability to maintain functional flexibility despite the stability of its structural architecture is critical for cognition and health.

Given the foundational role of structure-function flexibility in brain function, understanding how this relationship changes with aging is crucial for understanding cognitive decline and health deterioration in older adults. Although research has identified the multi-level impacts of aging on SC and FC, including white matter damage, myelin degradation of SC, and the breakdown of network modularity in FC, the exploration of the flexible and dynamic relationship between SC and FC remains in its early stages. Several studies have explored how SFC changes with age (Baum et al., 2020; Vandewouw, Hunt, Ziolkowski, & Taylor, 2021; Zamani Esfahlani, Faskowitz, Slack, Mišić, & Betzel, 2022; Z. Zhang et al., 2025), but few have focused specifically on aging. These studies report that aging is associated with widespread reductions in SFC, particularly in unimodal sensory and motor cortices, with the most pronounced declines observed in the visual and sensorimotor areas, likely due to the loss of white matter myelination (Vandewouw et al., 2021; Zamani Esfahlani et al., 2022). Abnormalities in SFC have also been observed in aging-related brain diseases such as Alzheimer’s disease (Cao et al., 2020; Dai et al., 2019; Sun et al., 2014), Parkinson’s disease (Zarkali et al., 2021), and stroke (J. Zhang et al., 2017).

Despite these findings, most research has treated SFC as a static feature, largely overlooking its dynamic properties. Over the past decade, it has become increasingly clear that brain function is inherently dynamic, with functional connectivity exhibiting time-varying fluctuations that are non-random and organized (Allen et al., 2012; Hutchison et al., 2013; Preti, Bolton, & Van De Ville, 2016). Given the relative stability of brain structure, functional activity must continuously adapt to external environments. Therefore, to fully understand the relationship between brain structure and function, it is essential to consider both the static and dynamic aspects of structure-function coupling. Some studies have already investigate the dynamic characteristics of structure-function coupling (Fotiadis et al., 2023; Liu et al., 2022; Preti & Van De Ville, 2024; Z. Zhang et al., 2024) and found SFC was moderated by cognitive demands (Griffa, Amico, Liégeois, Van De Ville, & Preti, 2022) and state of consciousness (drowsiness) (Preti & Van De Ville, 2024), and dynamic decoupling is particularly pronounced in brain networks that engage complex cognitive functions, such as attentional and memory tasks (Fotiadis et al., 2023; Liu et al., 2022). While much of this research has focused on younger adults, there is a notable gap in our understanding of how these dynamics evolve across the lifespan, particularly in middle-to-late adulthood, how they contribute to cognitive decline and changes in health status, and what genetic and environmental factors influence them.

To address these gaps, we combined traditional SFC methods with dynamic analysis techniques, leveraging multimodal neuroimaging data from 34,067 participants (ages 45–82) in the UK Biobank to calculate both static SFC and dynamic SFC (DSFC). Our goal was to investigate how both SFC and DSFC evolve across the adult lifespan and how these dynamics change with aging. Specifically, we aimed to answer three interrelated questions: (1) How do SFC and DSFC change with advancing age? (2) How do SFC and DSFC support cognitive function and influence health status? (3) What genetic and environmental factors modulate these trajectories? By integrating neuroimaging, genetic, and deep phenotypic data, our study provides a comprehensive examination of how structural and dynamic aspects of brain organization evolve during aging and how they relate to cognitive resilience and vulnerability.

## Results

### Static & Dynamic Structure-Function Coupling have different spatial pattern

To characterize the temporal variability of structure–function coupling (SFC), we computed dynamic structure–function coupling (DSFC) using a sliding-window approach. The window length was set to 30 TRs (0.735s×30 = 22.05s) with a step size of 5 TR. Within each window, a functional connectivity (FC) matrix was estimated using Pearson’s correlation across all brain regions. For a given brain region, the SFC at each time window was quantified as the Spearman rank correlation between its functional- and structural-connectivity profile (streamline number). Static SFC was defined as the mean SFC across all windows, whereas DSFC was defined as the standard deviation of SFC across windows, reflecting temporal variability in structure–function coupling. We divided it according to Yeo 7 network parcellation.

At the group level, we first averaged SFC and DSFC across individuals (Fig. 1A) and then organized the seven canonical functional networks into four quadrants based on their mean SFC and DSFC values (Fig. 1B). the visual and dorsal attention networks exhibited the highest static structure-function coupling (SFC), while the frontoparietal network (FPN) showed the highest dynamic structure-function coupling (DSFC). The visual network (VN) and dorsal attention network (DAN) tended to exhibit high SFC with low DSFC, while the limbic network (LN) showed low coupling and low dynamics. The frontoparietal network (FPN) was characterized by high SFC and high DSFC, while the default mode network (DMN), ventral attention network (VAN), and sensorimotor network (SMN) generally fell in between, exhibiting medium SFC and medium DSFC (Fig. 1A). This spatial organization was consistent across alternative structural connectivity metrics (e.g., mean fractional anisotropy/FA of white-matter tracts between regions, see Supplementary Materials). Furthermore, DSFC maintained individual-level variability across streamline-based and FA-based methods (at network-level, Pearson r=0.83∼0.97, see Supplementary Materials). These results demonstrated that both SFC and DSFC are robust and reliable measures, capturing meaningful individual differences in dynamic structure-function coupling.

**Figure 1.**
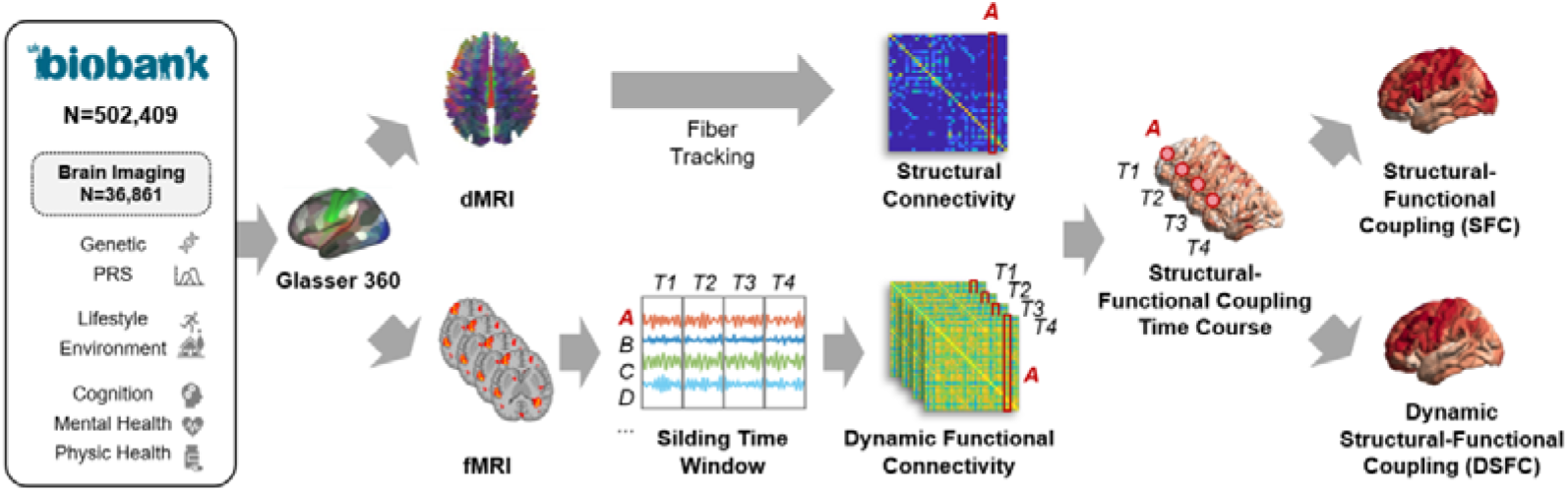
workflow of SFC and DSFC. Overview of multimodal data processing and structure-function coupling analysis. Data were obtained from the UK Biobank (N = 502,409; brain imaging after quality control, N = 34,067) including genetics, biomarkers, lifestyle, environment, cognition, mental health, and physical health. Structural connectivity was derived from diffusion MRI (dMRI) using fiber tracking and the Glasser 360 parcellation. Functional MRI (fMRI) data were processed with a sliding time window approach (window size=30 TRs, step size=5 TRs) to compute dynamic functional connectivity (T1, T2, T3, T4, …). For each time window, Spearman’s correlation was computed between structural and functional connectivity patterns across brain regions (e.g., brain region A shown in the figure) and the rest of the brain, serving as the structure-function coupling measure for that time window. Static structure-function coupling (SFC) was calculated by averaging across all time windows, while dynamic structure-function coupling (DSFC) was measured by computing the temporal standard deviation across time windows.

**Figure 2.**
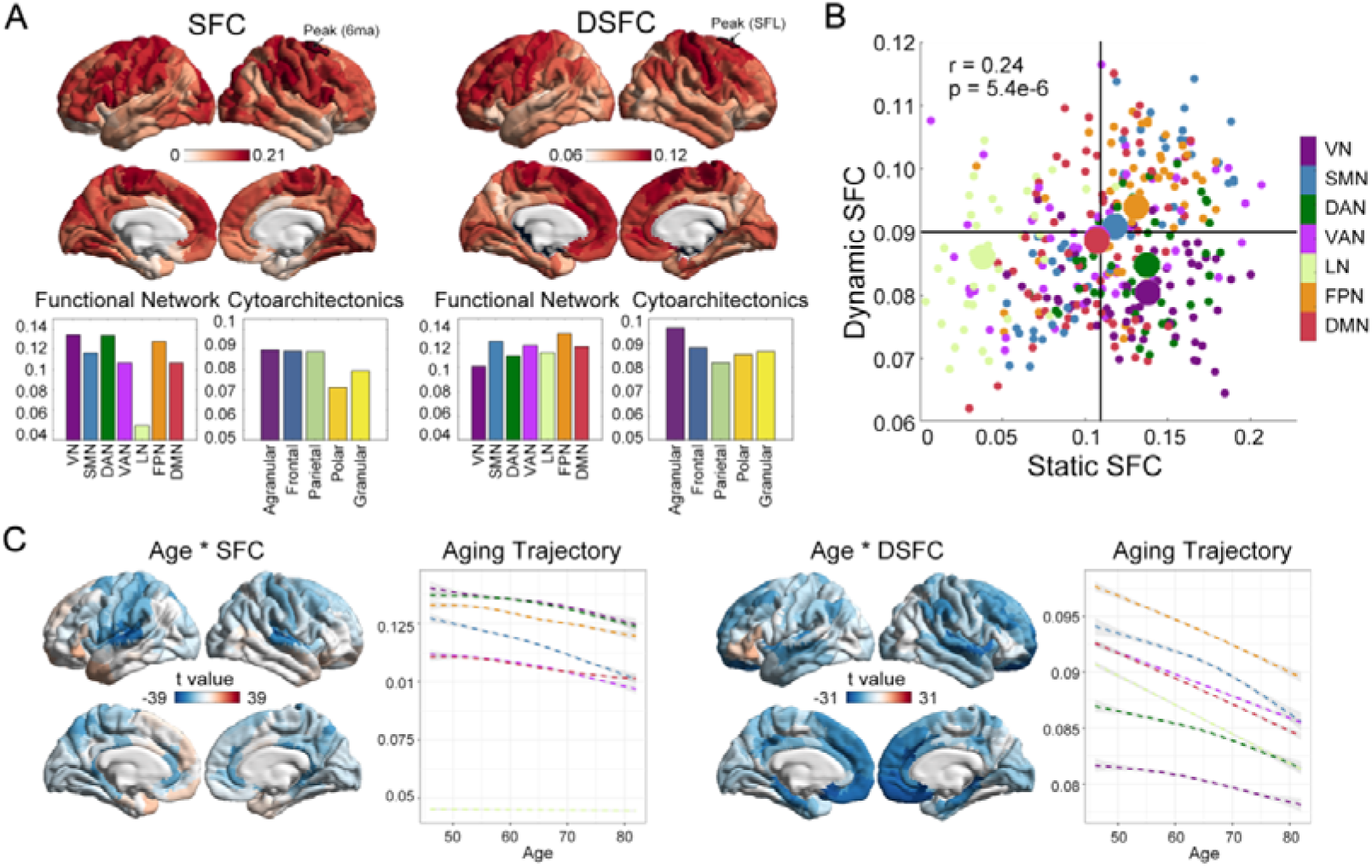
Spatial organization and aging trajectories of SFC and DSFC. (A) Cortical surface maps of group-averaged static SFC and dynamic DSFC. Functional networks are delineated according to the Yeo 7-network parcellation. (B) Scatter plot illustrating the regional relationship between SFC and DSFC. Each dot represents a brain region, color-coded by its functional network assignment. (C) Aging trajectories of SFC and DSFC. Left panels: Aging profiles of SFC showing significant declines primarily in sensorimotor systems. Right panels: Aging trajectories of DSFC highlighting pronounced reductions in higher-order networks (e.g., DMN, FPN). Statistical significance was assessed using Generalized Additive Mixed Models (GAMMs); t-values for the age effect are indicated for each network.

### SFC and DSFC show distinct aging effect

To examine the impact of aging on static and dynamic structure-function coupling (SFC and DSFC), we employed both generalized additive mixed models (GAMMs) and linear mixed models (LMMs). GAMMs were used to plot the aging trajectories of SFC and DSFC across seven networks. In contrast, LMMs were applied to assess the linear relationship between age and SFC/DSFC in each brain region. Our analysis revealed that both SFC and DSFC declined with age, though following distinct aging trajectories. The age-related decline in SFC was primarily confined to the sensorimotor networks (t = -35, p = 7.8e-264). In contrast, DSFC showed a much stronger aging effect, with the most significant reductions observed in higher-order networks, including DMN (t = -34.8, p = 7e-261), FPN (t = -27.1, p = 2.1e-159), and limbic network (t = -39.62, p < 5e-300). This pattern of decline was consistent across different structural connectivity measures, such as fractional anisotropy (FA) and streamline count (SM Figure 1). These findings suggest that aging selectively impairs the ability of higher-order brain regions to flexibly generate functional connectivity patterns, independent of their structural constraints (Fig. 1B).

### SFC and DSFC are associated with distinct cognitive and health phenotypes

To further explore the relationship between static and dynamic structure-function coupling (SFC and DSFC) and a broad range of phenotypic traits, we employed linear mixed-effects models (LMMs) to examine their associations with cognitive, mental health, and physical health measures from the UK Biobank. The results revealed distinct patterns of association, with significant differences in the link between SFC and DSFC across cognitive and health domains.

Notably, stronger SFC in higher-order cognitive networks, such as the FPN and DMN, was associated with a wide range of higher cognitive functions, including fluid intelligence (FPN, t=5.8, p=6.6e-9; DMN, t=7.32, p=2.6e-13), the number of puzzles solved correctly (FPN, t=7.79, p=7e-15; DMN, t=7.19, p=6.4e-13), and digit span working memory (FPN, t=4.53, p=5.9e-6; DMN, t=5.31, p=1e-7). Parallel to these findings, widespread cognitive networks—including the DAN, VAN, limbic network (LN), FPN, and DMN—showed stronger positive correlations with these cognitive functions for DSFC than for SFC. Specifically, this trend was evident in fluid intelligence (DAN, t=9.72, p=2.6e-22; VAN, t=9.14, p=6.9e-20; LN, t=9.4, p=5.9e-21; FPN, t=5.48, p=4.3e-8; DMN, t=6.81, p=1e-11), puzzles solved (DAN, t=7.33, p=2.4e-13; VAN, t=8.47, p=2.55e-17; LN, t=6.13, p=8.9e-10; FPN, t=5.68, p=1e-8; DMN, t=7.57, p=4e-14), and digit span working memory (DAN, t=6.92, p=4.63e-12; VAN, t=6.19, p=6e-10; LN, t=6.72, p=1.8e-11; FPN, t=4.64, p=3.47e-6; DMN, t=5.02, p=5.09e-7). Region-level association analyses further revealed that these correlations spanned large brain areas, including the temporal lobes, insula, inferior frontal gyrus, and temporo-parietal junction (Fig. 3B). These results highlighted the prominent role of DSFC in supporting cognitive abilities, suggesting that SFC and DSFC in higher-order networks contribute jointly to cognitive performance.

**Figure 3.**
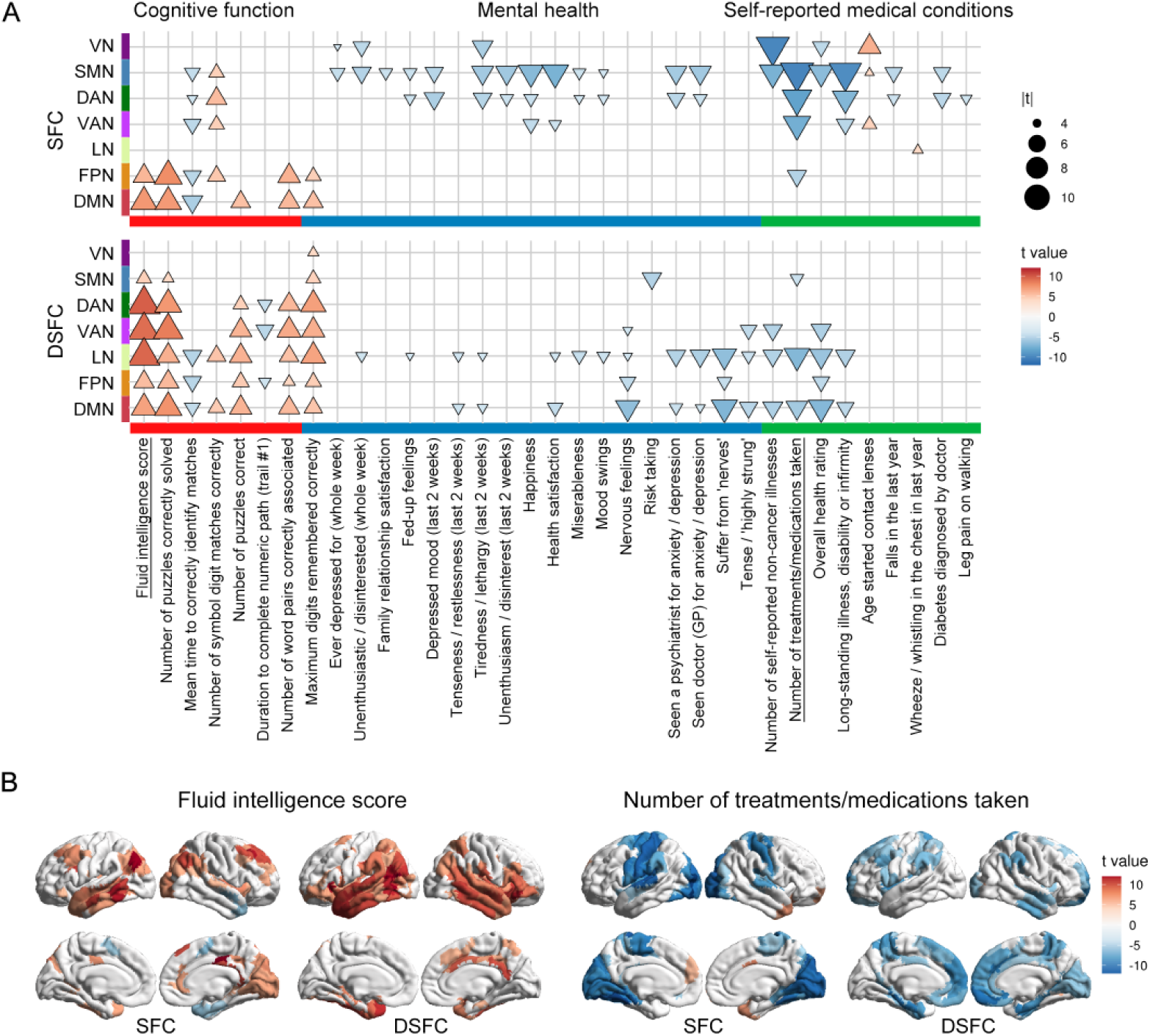
Associations of SFC and DSFC with cognitive performance and health conditions. (A) Heatmap showing the associations (t-values) of SFC (top) and DSFC (bottom) across various cognitive (green), mental health (red), and physical health (blue) phenotypes. Statistical significance was determined using Linear Mixed Models (LMMs), with age, sex, and head motion as covariates. (B) Cortical surface maps showing the spatial distribution of associations with fluid intelligence (positive association) and the number of medications taken (negative association). Red-yellow clusters indicate positive t-values, while blue-light blue clusters indicate negative t-values. Results are thresholded at p < 0.05 (FDR corrected).

In contrast, SFC and DSFC exhibited a clear spatial separation in their association pattern with physical/mental health factors. Significant associations were observed between SFC in lower-order networks, such as the SMN and VN, and various health-related variables, including the number of self-reported non-cancer illnesses (SMN, t=-9.99, p=1.75e-23; VN, t=-7.36, p=1.85e-13), the number of treatments/medications taken (SMN, t=-10.1, p=6e-24), frequency of unenthusiasm/disinterest in the last two weeks (SMN, t=-5.92, p=3.16e-9; VN, t=-4.81, p=1.49e-6), doctor visits for anxiety or depression (SMN, t=-5.86, p=4.43e-9; VN, t=-3.34, p=0.0008), overall health rating (SMN, t=-6.69, p=2.3e-11; VN, t=-4.8, p=1.53e-6), health satisfaction (SMN, t=-7.34, p=2.17e-13; VN, t=-3.19, p=0.0014), long-standing illness or disability (SMN, t=-9.65, p=5.49e-22; DAN, t=-7.04, p=1.92e-12), and age when glasses or contact lenses were first worn (VN, t=6.15, p=7.9e-10). Conversely, lower DSFC in higher-order networks, such as the DMN and LN, was linked to various health and affective symptoms, including overall health rating (DMN, t=-6.96, p=3.5e-12; LN, t=-6.14, p=8.3e-10), anxiety symptoms (DMN, t=-7.1, p=1.28e-12; LN, t=-6.52, p=6.92e-11), and the number of treatments/medications taken (DMN, t=-5.79, p=7e-9; LN, t=-6.99, p=2.69e-12) (Fig. 3B).

### Environmental and Lifestyle Modulators of SFC and DSFC

Similar LMMs were used to examine the associations between static and dynamic structure-function coupling (SFC and DSFC) and various environmental and lifestyle factors, including demographic variables, socioeconomic status, smoking, alcohol consumption, exercise habits, dietary habits, and early-life trauma. Overall, the associations between environmental and lifestyle factors and SFC/DSFC were less pronounced than those observed for cognitive abilities and health conditions (Fig. 4A). The most notable finding was that smoking was associated with a reduction in SFC within the DAN (t = -8.43, p = 3.72e-17) and FPN (t = -6.89, p = 5.74e-12) (Fig. 4B).

**Figure 4.**
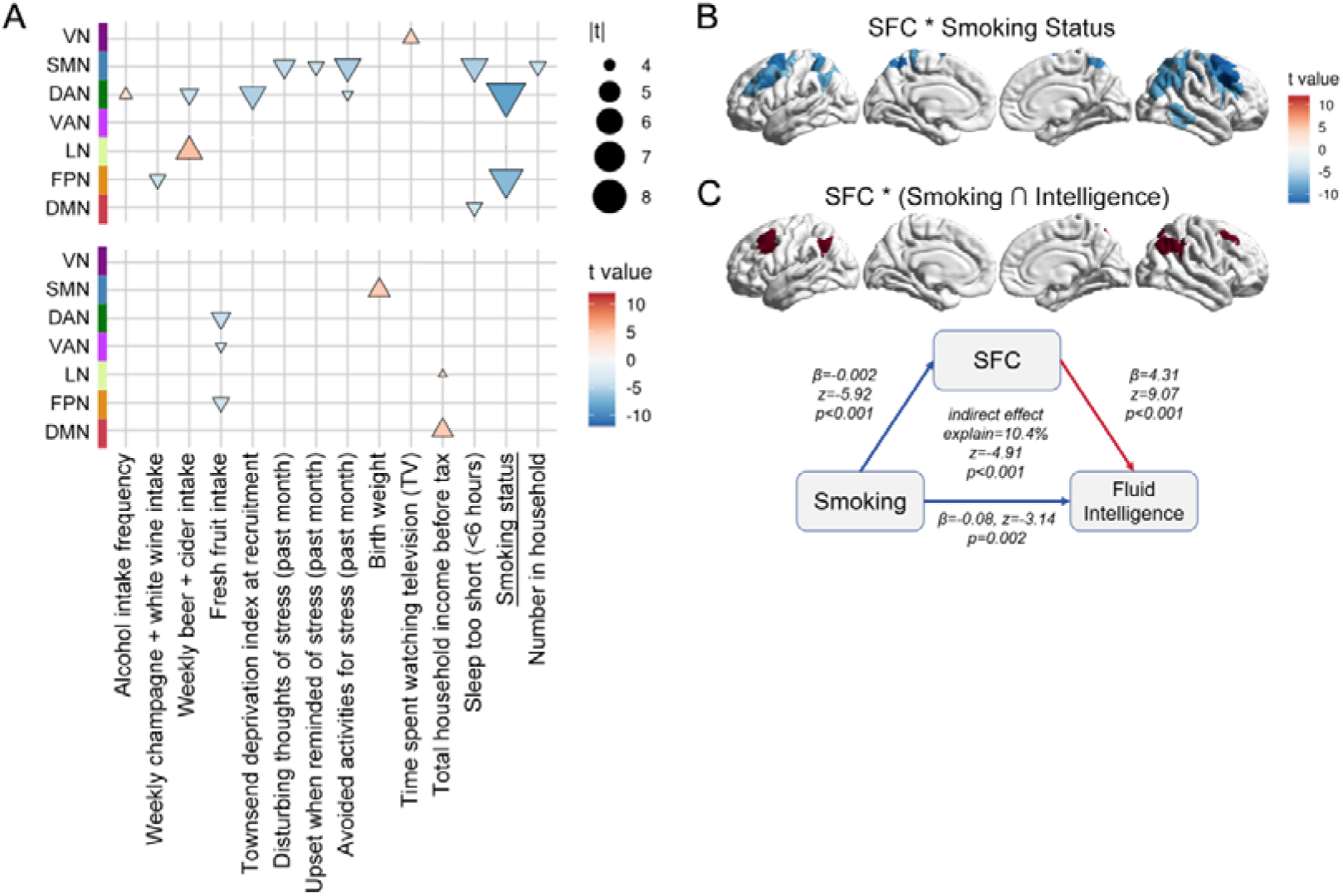
Environmental influences and mediation pathways. (A) Matrix plot illustrating associations between functional network coupling and lifestyle/demographic factors. Circle size and color intensity represent the magnitude and direction of the t-statistic, respectively. (B) Regional brain maps highlighting the significant impact of smoking on SFC and household income on DSFC. (C) Structural equation model (SEM) depicting the mediating role of SFC in the relationship between smoking and fluid intelligence. Path coefficients represent standardized betas (β). The indirect effect and its significance (p-value) were estimated using bootstrapping (10,000 iterations).

Using mediation analysis, we found that smoking could potentially reduce future cognitive performance by lowering SFC in the superior parietal lobule and frontal eye fields (FEF) (indirect effect explained = 10.4%, z = −4.91, p < 0.001) (Fig. 4C). For DSFC, environmental influences were weaker compared to those on SFC (Fig. 4A), with the most significant effects being increased DSFC in the DMN associated with higher household income (t = 4.72, p = 2.37e-6) and increased DSFC in the SMN related to birth weight (t = 4.73, p = 2.24e-6).

### Genetic Architecture of Structure-Function Coupling

Apart from the external influences of environment and lifestyle, we investigated whether individual differences in SFC and DSFC were also regulated by internal genetic factors. To explore the molecular genetic mechanisms underlying SFC and DSFC, we conducted network-level genome-wide association analyses (GWAS) to identify single nucleotide polymorphisms (SNPs) significantly associated with these measures. Our results revealed no genome-wide significant SNPs (p < 5e-8). These findings suggested that individual differences in both static and dynamic structure–function coupling were likely driven by the polygenic accumulation of numerous small-effect variants, rather than by any single locus of major effect.

### Polygenic Risk for Alzheimer’s Disease Selectively Affects Coupling Dynamics

Due to the limited explanatory power of individual SNPs (as shown in the GWAS results), we subsequently employed polygenic risk score (PRS) analysis to capture cumulative genetic effects. To investigate how genetic risk influences structure–function coupling and its dynamics, we examined the associations between SFC/DSFC and a comprehensive set of 87 PRSs, including 39 standard PRSs related to a broad range of physical, mental, and neurological disorders, as well as 51 enhanced PRSs associated with physiological quantitative traits (Thompson et al., 2022, 2024). Among these 87 disorders. We found that the PRS of Alzheimer’s disease (AD) showed the strongest negative association with DSFC, particularly in higher-order cortical networks such as the SMN (t = −4.46, p = 8.34e-6), VAN (t = −5.02, p = 5.31e-7), limbic network (t = −4.53, p = 5.81e-6), FPN (t = −5.49, p = 3.96e-8), and DMN (t = −4.78, p = 1.78e-6) (Fig. 5A&D). In contrast, only SFC in the VN showed significant associations with AD genetic risk (t = −4.73, p = 2.27e-6) and schizophrenia (t = −4.92, p = 8.79e-7). These results suggest that dynamic coupling may reflect a genetically sensitive aspect of network adaptability that precedes overt structural decline.

**Figure 5.**
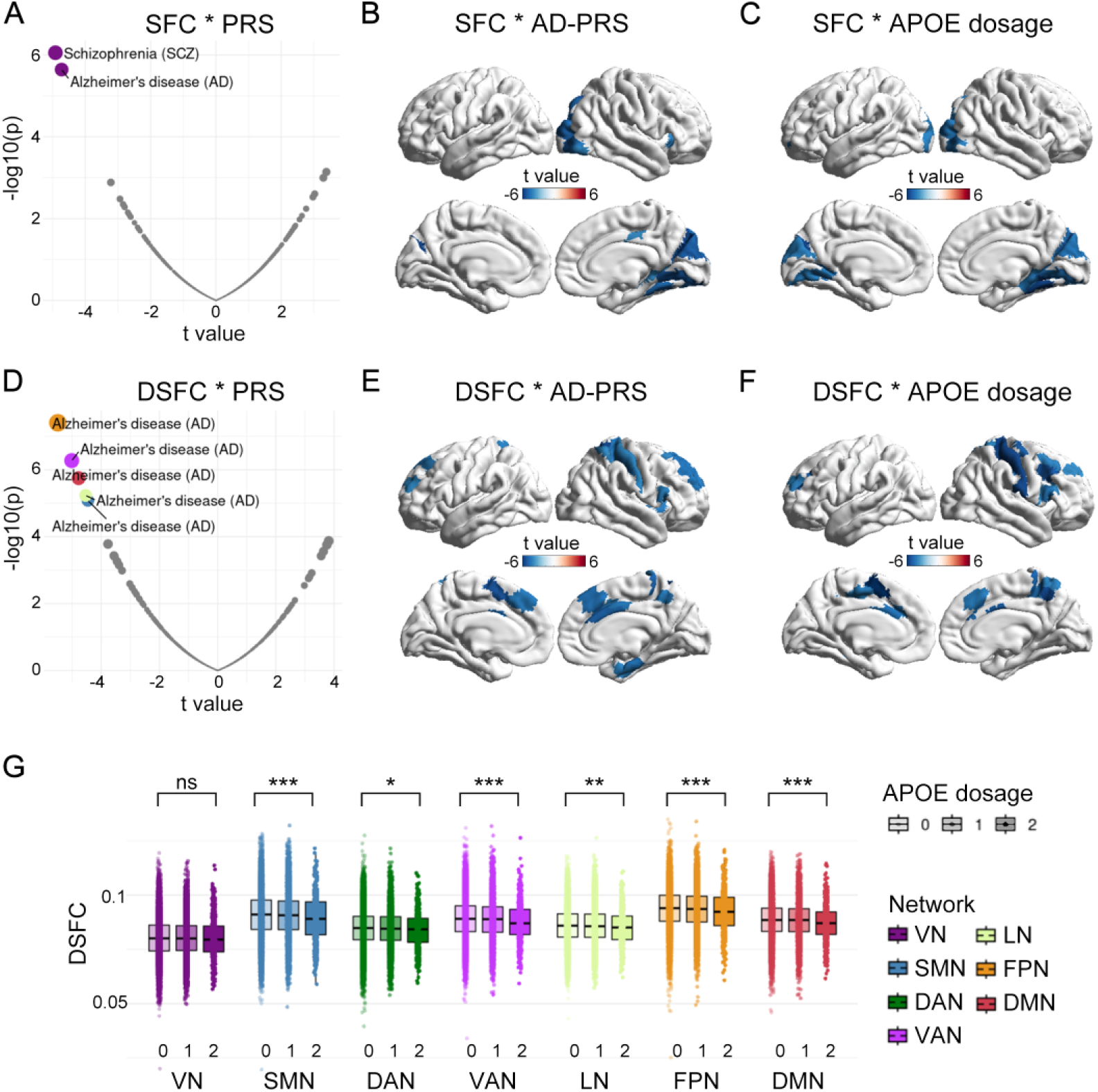
Associations of SFC and DSFC with polygenic risk scores and APOE ε4 dosage. (A, D) Volcano plots (or Manhattan-style plots) showing the association between network-level (A) SFC and (D) DSFC with 87 polygenic risk scores (PRSs). The y-axis represents -log10(p-values) and the x-axis represents t-statistics. (B, E) Cortical surface maps illustrating the spatial distribution of associations (t-values) between (B) SFC and (E) DSFC with the PRS for Alzheimer’s disease (AD-PRS). (C, F) Brain maps showing the regional associations of (C) SFC and (F) DSFC with APOE ε4 allele dosage (0, 1, or 2). (G) Network-specific effects of APOE ε4 dosage. Box plots illustrate the distribution of coupling strength across ε4 dosage groups for each functional network (VN, SMN, DAN, VAN, LN, FPN, DMN). Inter-group differences were assessed using the Kruskal-Wallis test, with significance levels denoted as: *, p < 0.05, **, p < 0.01$, ***, p < 0.001.

**Figure 5.**
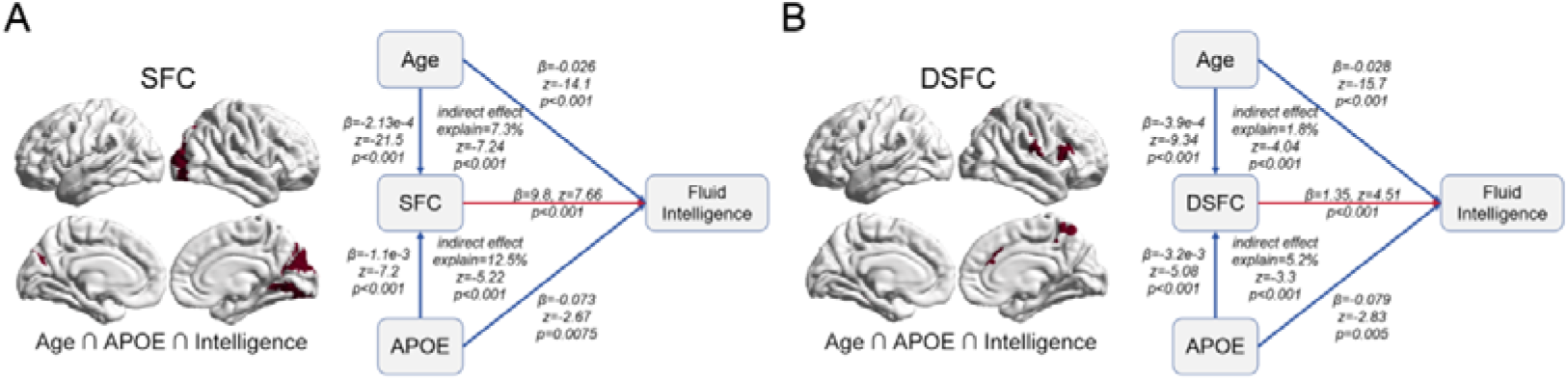
Mediation pathways linking aging and genetic risk to cognitive decline. (A) Static SFC mediation: Structural equation model (SEM) illustrating the mediating role of SFC (specifically within occipital visual cortex) in the relationship between age, APOE ε4 dosage, and fluid intelligence. (B) Dynamic SFC mediation: Parallel mediation model for DSFC (specifically within the ventral attention/higher-order networks). In both models, standardized path coefficients (β) and their significance levels are provided for direct and indirect effects. Intersection brain regions used for calculating mediator values are highlighted in red on the surface maps. All indirect effects were tested using bootstrapping with 5,000 iterations.

### Impact of APOE ε4 Dosage on SFC and DSFC Trajectories

Given the specific strong correlation between AD polygenic risk and DSFC, we sought to further identify the specific genetic drivers of this association. Based on the most recent meta-analysis (Andrews et al., 2023), we extracted 81 SNPs from 21 candidate loci. Using linear mixed models (LMMs) controlling for standard covariates and 20 genetic principal components (PCs), we identified a specific association between DSFC and SNPs located within the Apolipoprotein E (APOE) gene.

We further quantified APOE ε4 dosage and examined its relationship with SFC and DSFC across large-scale brain networks, while also testing whether age moderated these associations. We observed that individuals carrying a higher ε4 allele dosage exhibited lower SFC in the VN (t = −5.15, p = 2.69e-7), particularly in the ventral visual complex (VVC; t = −5.56, p = 2.7e-8). Simultaneously, higher ε4 dosage was associated with lower DSFC across multiple networks, including the SMN (t = −4.5, p = 6.67e-6), VAN (t = −4.83, p = 1.34e-6), LN (t = −4.72, p = 2.41e-6), FPN (t = −5.24, p = 1.65e-7), and DMN (t = -3.95, p = 7.74e-5), with the most pronounced reduction in the supplementary and cingulate eye field (SCEF; t = −5.98, p = 2.28e-9). Notably, this DSFC reduction was already evident in middle-aged individuals, suggesting that ε4-related dynamic impairment emerges relatively early. In contrast, the association between ε4 dosage and SFC was age-dependent: during midlife, SFC showed little relationship with ε4 dosage, whereas in older adults, higher dosage was significantly associated with reduced SFC. These findings suggest that ε4 allele dosage begins to affect DSFC as early as midlife, indicating that DSFC may serve as a more sensitive early biomarker of neurodegeneration than static SFC.

### SFC and DSFC Mediate the Effects of Aging and Genetic Risk on Cognitive Function

To investigate whether SFC and DSFC acted as mediators linking genetic factors (APOE ε4 dosage) and age to cognitive outcomes (fluid intelligence), we conducted a multiple mediation analysis. We averaged the SFC or DSFC values in brain regions that were significantly associated with age, ε4 allele dosage, and cognitive performance (the intersection regions) and used these averages as mediators in the model.

Our results revealed significant mediation effects: SFC in the occipital lobe, along with DSFC in the inferior frontal gyrus and dorsal anterior cingulate cortex (primarily within the VAN), significantly mediated the relationship between age and higher ε4 allele dosage with fluid intelligence scores. Specifically, the mediation effects were as follows: For SFC, the indirect effect of age was significant (β = -2.1e-3, 95% CI [-2.67e-3, -1.5e-3], z = -7.24, p = 4.39e-13), explaining 7% of the total variance; the indirect effect of APOE ε4 dosage was also significant (β = -1.06e-2, 95% CI [-1.48e-2, -6.81e-3], z = −5.22, p = 1.8e-7), explaining 12%.

For DSFC, the indirect effect of age was β = −5.3e-4 (95% CI [-8e-4, -3e-4], z = −4.04, p = 5.24e-5), explaining 1.8% of the variance; the indirect effect of APOE ε4 dosage was β = −4.37e-3 (95% CI [-7e-3, -2e-3], z = -3.3, p = 9.73e-4), explaining 5.2%. These findings suggested that SFC and DSFC played crucial roles in mediating genetic and age-related influences on cognitive outcomes, highlighting their potential as biomarkers for understanding the mechanisms of cognitive aging.

## Discussion

In this study, leveraging an extensive neuroimaging dataset from over 34,000 participants, we characterized the complementary roles of static and dynamic structure–function coupling (SFC and DSFC) in aging and cognitive preservation. Our findings revealed that both SFC in sensorimotor systems and DSFC in higher-order networks declined significantly with age. Crucially, we demonstrated that while SFC and DSFC in higher-order networks supported cognitive performance, their disruptions—specifically within sensory-motor SFC and higher-order DSFC—linked to heightened mental and physical health burdens. By integrating genetic and environmental evidence, we showed that aging and Alzheimer’s disease-related risk factors, notably the APOE ε4 dosage, modulated these coupling patterns to influence longitudinal cognitive outcomes. We propose a conceptual framework where SFC reflects the stability of the brain’s core architecture, while DSFC captures the functional flexibility required to switch between coupling and decoupling states. These two mechanisms likely serve as synergistic pillars of higher-order cognition. Consequently, preserving the integrity of both network stability (SFC) and dynamic adaptability (DSFC) may be essential for mitigating age-related cognitive decline and maintaining brain health in older adults.

### Complementary Roles of Static and Dynamic Structure-Function Coupling in Cognitive Function and Health

We found that both SFC and DSFC in higher-order cognitive networks are positively correlated with cognitive performance, including fluid intelligence, problem-solving ability, and working memory. This aligns with prior research showing that SFC predicts various cognitive aspects, such as working memory (Baum et al., 2020), cognitive flexibility (Medaglia et al., 2018), fluid intelligence (Griffa et al., 2022) and composite cognitive scores (Gu et al., 2021). Our study extends this understanding by revealing that dynamic structure-function coupling, particularly DSFC, has a stronger association with cognitive performance than static SFC, emphasizing the importance of flexible network reconfiguration in maintaining cognitive abilities (Braun et al., 2015; Ju & Bassett, 2020; Shine, 2019). The association between DSFC in the temporal cortex and cognitive performance is especially pronounced (Fig. 3B). The temporal cortex is crucial for conceptual categorization, semantic processing, language function, and associative memory (Braunsdorf et al., 2021; Jackson, Bajada, Rice, Cloutman, & Lambon Ralph, 2018). A decline in its ability to decouple from structural constraints may impair integration with other networks, hindering the transfer of semantic information and memory to working memory, thus impacting cognition. Overall, our findings highlight the complementary roles of both SFC and DSFC in supporting cognitive function, with structural and functional coherence, alongside flexibility, being key to human intelligence.

In the context of health, we observed distinct spatial patterns in the effects of mental and physical health burdens on SFC and DSFC: poorer health is associated with lower SFC in sensorimotor regions and higher DSFC in higher-order cognitive networks. Sensory and motor networks exhibit the highest levels of SFC across the brain (Dong et al., 2024; Fotiadis et al., 2023; Gu et al., 2021), with the highest cortical myelination (Glasser, Goyal, Preuss, Raichle, & Van Essen, 2014), and their functional activity is strongly constrained by white matter connectivity, supporting the high precision and robustness required for sensory-motor functions. Physical and mental health problems (such as depression and anxiety) and associated chronic stress activate immune responses, triggering neuroinflammation that impairs oligodendrocyte function, leading to myelin abnormalities, which might further reducing SFC (Antontseva, Bondar, Reshetnikov, & Merkulova, 2020; Gorlova et al., 2023; Knowles, Batra, Xu, & Monje, 2022; Murayama, Cai, Nakamura, & Hashimoto, 2025). Additionally, poor health may impair long-range white matter connectivity (de Lange et al., 2019; Marebwa et al., 2018; Meijer, Steenwijk, Douw, Schoonheim, & Geurts, 2020), compromising the brain’s ability to integrate across brain modules (Betzel & Bassett, 2018), preventing higher cognitive networks from effectively decoupling from structural constraints, and ultimately reducing cognitive flexibility.

### Aging, Cognitive Decline, and Static and Dynamic Structure-Function Coupling

Our results show that static and dynamic structure-function coupling follow distinct aging trajectories, with SFC in sensorimotor regions and DSFC in higher-order cognitive networks decreasing with age. The finding regarding SFC aging trajectory is consistent with previous studies (Zamani Esfahlani et al., 2022), which demonstrated a decline in global structure-function coupling with aging, driven by reductions in sensorimotor network coupling. SFC have been linked to myelination (Fotiadis et al., 2023), and the degeneration of SFC may be associated with myelin degradation in older adults, along with their decreased sensorimotor function (Fleischman et al., 2015; Gong et al., 2025; Wakefield et al., 2010; Yoshimura et al., 2020). Our findings also align with results showing reduced SFC in these regions among individuals with higher stress levels and poorer physical and mental health, who may have elevated inflammation, contributing to the disruption of structure-function networks related to bodily symptoms.

Our findings corroborate previous research on static SFC and extend it to the study of dynamic SFC, revealing that, with aging, the ability of higher-order cognitive networks to switch between coupling and decoupling states declines. Our mediation analysis further demonstrates that aging influences fluid intelligence performance by modulating the level of DSFC, which further strengthens the chain of relationships between aging, the decline in the dynamic switching capability between coupling and uncoupling in higher-order networks, and cognitive decline. Prior graph-theoretical analyses have shown that these regions exhibit higher participation coefficients (Kulkarni & Bassett, 2025; Pedersen, Omidvarnia, Shine, Jackson, & Zalesky, 2020; Power, Schlaggar, Lessov-Schlaggar, & Petersen, 2013), indicating functional integrating across multiple functional modules. The higher-order networks also occupy top positions in the global brain functional gradient and play a critical role in integrating information across spatial and temporal scales (Huntenburg, Bazin, & Margulies, 2018; Margulies et al., 2016; Vázquez-Rodríguez et al., 2019). This suggests that the flexibility of these networks reflects the brain’s ability to decouple from structural constraints, enabling whole-brain integration and the recruitment of global cognitive resources for complex tasks. A decline in this flexibility indicates reduced brain adaptability, contributing to cognitive decline in older adults or patients.

We observed that the dynamic characteristics of later-developing higher-order cognitive networks appear to enter a decline phase earlier, supporting the classic "last-in, first-out" hypothesis in developmental neuroscience (Whalley, 2015). Early SFC development studies have shown an opposite trend, with SFC in primary sensory systems declining throughout life, while SFC in higher-order cognitive networks increases during childhood and adolescence, stabilizing in early adulthood (Baum et al., 2020; Zamani Esfahlani et al., 2022; Z. Zhang et al., 2025). Our study adds a dynamic perspective to this framework, highlighting the significance of dynamic decoupling and coupling strength in assessing different aspects of the structure-function relationship and its development during the lifespan.

### Biological Mechanisms and Environmental Influence Factors of Static and Dynamic Coupling

We found that APOE ε4 dosage negative associate with SFC in visual cortex and DSFC in SMN and higher-order cognitive networks (such as FPN and VAN). APOE, a key protein involved in lipid transport and the clearance of abnormal proteins in the brain, is primarily synthesized by astrocytes and microglia. APOE ε4 impairs the brain’s ability to clear low-density lipoprotein cholesterol (LDL-C), increasing the risk of cardiovascular disease and Alzheimer’s disease (AD). Previous studies have shown that APOE ε4 carriers exhibit reduced network efficiency (V. Heise, Filippini, Ebmeier, & Mackay, 2011) and lower white matter integrity (Verena Heise et al., 2024). These changes are also associated with a significant decrease in the efficiency of the medial temporal lobe functional networks in cognitively normal elderly individuals (Chen et al., 2015). We hypothesize that APOE ε4 may affect structural-functional coupling by reducing white matter efficiency, weakening visual network robustness, and impairing higher cognitive regions’ communication. Our GWAS analysis did not reveal significant genetic associations, which contrasts with twin studies showing higher heritability of SFC in visual, subcortical, cerebellar, and brainstem (Valk et al., 2022). This discrepancy may stem from differences in methodology, as GWAS typically reports lower heritability than twin studies (Owen & Williams, 2021). GWAS generally detects smaller effects from individual SNPs, which can only be identified by aggregating multiple SNPs using approaches like polygenic risk scores (PRS). Additionally, our sample had a higher mean age compared to the HCP sample used in previous studies (around 30 years), which may have reduced the observed genetic effects.

We also identified the impact of environmental factors such as smoking habits, birth weight, and household income on SFC and DSFC. Smoking induces oxidative stress, increases inflammation, and may thus affect the white matter quality in higher-order brain regions (Brody et al., 2017; Gray et al., 2020; Yu et al., 2015). Previous studies have confirmed the effects of smoking on higher-order brain areas. Low birth weight, which indicates developmental or nutritional deficiencies, may affect the white matter development in sensorimotor networks, leading to abnormalities in white matter microstructure and reduced integrity (Alex et al., 2024; Eikenes et al., 2012; Neumane et al., 2022). Low household income is associated with chronic stress, which has been shown to accelerate brain aging and disrupt dynamic integration in higher-order brain regions (Blair & Raver, 2016; Farah, 2018; Krueger et al., 2025). Our findings provide insights into how environmental factors shape the structure-function relationships of the brain, offering a basis for interventions targeting environmental changes and lifestyle modifications to enhance cognitive function and health.

In addition to its role in aging, our findings have significant implications for understanding cognitive decline in the context of neurodegenerative diseases. Previous studies on SFC changes in Alzheimer’s disease (AD) have yielded inconsistent results, showing both increases and decreases (Cao et al., 2020; Dai et al., 2019; Sun et al., 2014), likely due to small sample sizes (the total sample size of all three studies was less than 100). Our research provides empirical evidence that clarifies the association between AD genetic risk and SFC and DSFC using an exceptionally large sample. Future studies may identify static and dynamic coupling decline as an early biomarker for aging and neurodegenerative diseases.

## Conclusion

In conclusion, our results emphasize the importance of dynamic structure-function coupling as a key mechanism for maintaining cognitive resilience in aging. While static SFC supports network stability, dynamic SFC enables the brain to adapt to changing demands, facilitating the flexibility necessary for cognitive performance. Disruptions in static and dynamic coupling, whether due to aging, genetic risk, or lifestyle factors, may contribute to cognitive decline, healthy risk and neurodegenerative diseases. Further research into the molecular and neurobiological mechanisms underlying these processes will be crucial for developing strategies to protect cognitive function in aging and disease.

## Limitations

Several limitations should be noted. First, this study is an association analysis rather than causal analysis, which limits our ability to infer causal relationships or track individual changes in structure–function coupling. Additionally, our GWAS analysis did not reveal significant genetic associations, likely because both SFC and DSFC are influenced by the cumulative effects of multiple SNPs. Such weak associations also make it impossible to perform Mendelian randomization using genetic variants as instrumental variables to investigate causal effects. Third, DSFC was estimated from fMRI signals, which are indirect measures of neural activity. Future research should combine neural electrophysiological activity to more directly measure functional networks. Finally, the main participants in this study were of British descent, which may limit the generalizability of the findings. Further studies are needed to validate and extend these results in more diverse cohorts.

## Methods

### Samples and participants

The UK Biobank (UKB) is a prospective cohort study involving ∼500,000 UK participants aged 40 to 69 years (Sudlow et al., 2015). Participant data encompass genomic and imaging data, electronic health record linkages, biomarkers, as well as physical and anthropometric measurements. Detailed information can be found at https://biobank.ndph.ox.ac.uk/showcase/. The UKB includes four instances (Instance 0 to 3). About 40,000 individuals (aged 44 to 82 years) underwent MRI scanning at Instance 2, which was collected during a follow-up visit approximately 5 to 9 years after baseline (Instance 0). Ethical approval for the UK Biobank study was granted by the Human Biology Research Ethics Committee at the University of Cambridge, and informed consent was obtained from all participants. This study utilized data from the UK Biobank under project number 19542.

### MRI data acquisition and Preprocessing

All UKB brain imaging data (connectome of Glasser parcellation, ID 31022; functional time series of Glasser parcellation, ID 31016) were acquired using 3T Siemens Skyra scanners with a standard 32-channel head coil. Further details about data and preprocessing are available in the online documentation: http://biobank.ctsu.ox.ac.uk/crystal/refer.cgi?id=2367 and http://biobank.ctsu.ox.ac.uk/crystal/refer.cgi?id=1977 (Alfaro-Almagro et al., 2018; Miller et al., 2016) and related article (Mansour L, Di Biase, Smith, Zalesky, & Seguin, 2023). The T1-weighted structural brain images were acquired using a 3D MPRAGE acquisition at 1 mm isotropic resolution with a 256 mm superior-inferior field of view.

Resting-state BOLD data were acquired with a multi-band gradient echo EPI sequence, with an acquisition time of 6 min (TR=0.735s), for a total of 490 volumes (Miller et al., 2016). Preprocessing steps (Alfaro-Almagro et al., 2018) consisted of the FSL MELODIC pipeline (Jenkinson, Beckmann Cf Fau - Behrens, Behrens Te Fau - Woolrich, Woolrich Mw Fau - Smith, & Smith). In addition to UKB quality control, participants with excessive head movement (average framewise displacement > 0.2mm, ID 25741) during scans were excluded.

Diffusion-weighted MRI (dMRI) data were acquired using a multi-band spin echo EPI sequence with a 7-minute acquisition time. The data were acquired across 100 unique diffusion sensitization directions, evenly distributed over two shells (b-values: 1000, 2000 s/mm^3^), along with 5 b=0 volumes. The spatial resolution was 2 mm isotropic voxels (MB=3, no in-plane acceleration, TE/TR=92/3600 ms, partial Fourier=6/8, conventional fat saturation). Additionally, 3 b=0 volumes were collected with reversed phase encoding to estimate susceptibility fields. Preprocessing including corrections for eddy currents, head motion, and gradient distortions (Alfaro-Almagro et al., 2018). Whole-brain tractography was estimated using *MRtrix3* (Tournier et al., 2019).

### Age effect analysis

We used linear mixed models (using the *fitlme* function in MATLAB R2018b) to detect aging effect of SFC and DSFC based on age at the time of brain scan (ID 21003, age at Instance 2). Sex (ID 31), race (ID 21000), body mass index (BMI, ID 21001), head movement (ID 25741), signal-to-noise ratio of resting state fMRI (SNR, ID 25744), and total intracranial volume (TIV, ID 26521) were used as fixed effects, and centre (ID 54) as random effects. Bonferroni correction was used to control I-type error. For regional nodal degree, multiple comparisons were corrected across 360 brain regions. At the network level, averaged SFC and DSFC were assessed across 7 network modules of Yeo 7 networks (Yeo et al., 2011). Generalized Additive Models (GAMs) was performed using the *gamm* function from the *mgcv* package in R (version 4.0), which were used to characterize age-related changes in the global redundant and synergistic components across the whole brain.

### Association study

We used linear mixed model to examine the associations between whole-brain averaged synergistic and redundant components and three categories of phenotypes: cognitive function (Category 100026), mental health (Category 100060), and self-reported medical conditions (Category 1003). To ensure consistency with the timing of brain imaging, we only included data from Instance 2. The analysis controlled for the same covariates used in the age-related analyses, including age itself as a fixed-effect covariate. We reported the associations of these measures with both regional- and network-level SFC/DSFC, with multiple comparisons corrected as previously described (360 brain regions and 7 measures of network-averaged SFC/DSFC).

Effect sizes were calculated based on the statistics provided by the linear mixed models (LMM). Since LMM do not directly yield *Cohen’s f^2^*and *Cohen’s d*, we adapted the methods: we used the regression coefficient (β) divided by the standard deviation of residuals to estimate *Cohen’s d*, and used the change in *R^2^* between a full and a reduced model (excluding the predictor of interest) to estimate *Cohen’s f^2^*. These effect sizes do not strictly conform to the standard definitions of *Cohen’s f^2^* and *Cohen’s d*. Therefore, the reported values are for reference only and are not presented in the main text.

We further examined the associations between redundancy/synergy and a broad range of environmental and lifestyle factors, using the same linear mixed-effects modeling as described above. Importantly, 63 environmental and lifestyle measures were derived from the UK Biobank baseline assessment (Instance 0), which temporally preceded brain imaging (Instance 2), thus ensuring temporal precedence of exposures relative to outcomes. The factors spanned multiple domains, including socioeconomic status (SES, e.g., household income, years of education), physical activity, social support (e.g., live alone, frequency of family/friend visits), sleep (e.g., sleep duration, insomnia), diet, alcohol use, smoking status, early life factors and traumatic childhood/adulthood experiences). For the analysis of environmental factors and lifestyle, since the measurements were taken prior to brain imaging, the time difference between the two measurements was also included as an additional covariate in the analysis.

Detailed variable IDs and descriptions are provided in Supplementary Table 1.

### Genetic analysis

Genotype data were available for all 500,000 participants in the UK Biobank cohort. Detailed genotyping and quality-control procedures for the UK Biobank are available in a previous publication (Bycroft et al., 2018) and online documentation: https://biobank.ctsu.ox.ac.uk/crystal/label.cgi?id=263. We excluded single-nucleotide polymorphisms (SNPs) with call rates <95%, minor allele frequency <0.1% or deviation from the Hardy–Weinberg equilibrium with P <5e-8. After the quality-control procedures, we obtained a total of 8,608,580 SNPs. We conducted genome-wide association analysis (GWAS) to investigate the relationship between genotype and 7 network-level SFC/DSFC. The GWAS was performed using PLINK 2.0 (Chang et al., 2015), adjusting for sex, age, BMI, head motion, SNR, TIV, centre, and the top 10 ancestry principal components.

### Polygenic Risk Scores (PRSs)

Polygenic risk scores (PRSs) were obtained from the UK Biobank’s curated PRS dataset (Category 2405)(Thompson et al., 2022, 2024), developed by Genomics PLC under UK Biobank project 9659. Version 2 of the dataset (released May 2024) was used in the present study. The PRSs were derived either entirely from external genome-wide association studies (the Standard PRS set) or from a combination of external and internal UK Biobank data (the Enhanced PRS set). Each PRS represents a weighted sum of allelic dosages across genome-wide variants, where weights correspond to effect sizes from discovery GWAS summary statistics. We included both Standard and Enhanced PRSs covering a wide range of complex traits and diseases, such as Alzheimer’s disease, schizophrenia, type 2 diabetes, body mass index (BMI), blood lipid profiles, and glycated haemoglobin (HbA1c). PRS values were standardized (z-scored) across participants prior to analysis. Associations between SFC/DSFC and PRSs were tested using linear mixed-effects models controlling for age, sex, BMI, head motion, SNR, TIV, genotype measurement batch (ID 22000), and the first 10 genetic principal components (ID 22009).

### APOE Genotyping

To determine APOE genotypes for all participants, we extracted the two defining single-nucleotide polymorphisms (SNPs) of the APOE locus—rs429358 (Chr19:45411941, GRCh37; reference allele = T, alternate allele = C) and rs7412 (Chr19:45412079, GRCh37; reference allele = C, alternate allele = T)—from the imputed UK Biobank genetic dataset. These two SNPs jointly encode the ε2/ε3/ε4 alleles of the APOE gene, which are determined by the amino acid substitutions at positions 112 and 158 of the apolipoprotein E protein. Genotype extraction was performed using PLINK 2.0 (build 20241124). Individuals were assigned APOE ε4 dosage according to the dosage of the alternate alleles at the two SNPs, following the canonical haplotype definition: ε2: rs429358 = T, rs7412 = T; ε3: rs429358 = T, rs7412 = C; ε4: rs429358 = C, rs7412 = C. In practice, we coded the count of the rs429358 C allele as the APOE ε4 dosage (0, 1, or 2), which captures the number of ε4 alleles carried by each individual. Associations between SFC/DSFC and PRSs were tested using linear mixed-effects models controlling for age, sex, BMI, head motion, SNR, TIV, genotype measurement batch, and the first 10 genetic principal components. Kruskal-Wallis Rank Sum Test (using *kruskal.test* function in R package *stats*) was employed to assess differences between the APOE ε4 dosage groups.

### Mediation Analysis

Mediation analyses were conducted using the *lavaan* package in R v4.0. We used genetic factors (APOE ε4 dosage) and age as independent variables, and cognitive outcomes (fluid intelligence) as dependent variables. SFC/DSFC in brain regions both significantly associated with age, ε4 allele dosage, and cognitive performance (intersection regions) was averaged as mediators in the model. Before performing the mediation analysis, fixed effects for sex, ethnicity, BMI, head motion, SNR and TIV, as well as the random effect of center, were removed using a linear mixed model (LMM). The residuals were then retained for input into the mediation model. For the mediation analysis of smoking-SFC-intelligence, we also included age as an additional covariate.

## Data availability

The data used in the present study are available from UK Biobank with restrictions applied. Data were used under licence and are thus not publicly available. Researchers can apply for access to the UK Biobank data via the Access Management System (https://www.ukbiobank.ac.uk/enable-your-research/apply-for-access). GWAS summary data for redundancy and synergy can be downloaded from the figshare (https://figshare.com/account/articles/30548912). European ancestry reference data from the 1000 Genomes Project can be found via https://github.com/getian107/PRScsx?tab=readme-ov-file.

## Code availability

For the analyses conducted in MATLAB 2025a and R (version 4.2.3). PLINK 2.0 was used to perform GWAS. The primary code used in this study has been made publicly accessible through the GitHub repository (https://github.com/XinRanWu/UKB_DynamicSFC).

## Acknowledgements

This study used the UK Biobank Resource under application number 19542. We thank all participants and researchers from the UK Biobank. J.Z. was supported by Science and Technology Innovation 2030 - Brain Science and Brain-Inspired Intelligence Project (STI2030-Major Projects 2021ZD0200204), and the State Key Laboratory of Neurobiology and Frontiers Center for Brain Science of Ministry of Education, Fudan University. This work was supported by the National Natural Science Foundation of China (no. 32500995 to X.W.). This work was partly supported by a grant from the National Key Research and Development Program of China (no. 2019YFA0709502 to J.F.), a grant from Shanghai Municipal Science and Technology Major Project (no. 2018SHZDZX01 to J.F.), Shanghai Centre for Brain Science and Brain-Inspired Technology and a grant from the 111 Project (no. B18015 to J.F.). The funders had no role in study design, data collection and analysis, decision to publish or preparation of the manuscript.

## Contributions

X.W., J.Z. conceived and designed the experiment. X.W. did the analyses with support from XXX and J.Z. X.W. drafted the paper with contributions from XXX and XXX and comments from all other authors. X.W. and XXX contributed to the visualization of data. X.W., S.P., XXX and J.Z. contributed to the interpretation of results. J.Z. and J.F. had full access to all the data in the study and took responsibility for the integrity of the data and the accuracy of the data analysis. All authors read and approved the final paper.

## Corresponding authors

Correspondence to Jie Zhang.

