## Supplemental Materials for "Disrupted dynamics of brain structure–function coupling link genetic risk of Alzheimer’s Disease and aging to cognitive decline in 34,067 adults"

**Supplemental Methods**

*MRI data acquisition and Preprocessing*

All UKB brain imaging data (connectome of Glasser parcellation, ID 31022; functional time series of Glasser parcellation, ID 31016) were acquired using 3T Siemens Skyra scanners with a standard 32-channel head coil. Further details about data and preprocessing are available in the online documentation: <http://biobank.ctsu.ox.ac.uk/crystal/refer.cgi?id=2367> and <http://biobank.ctsu.ox.ac.uk/crystal/refer.cgi?id=1977> (Alfaro-Almagro et al., 2018; Miller et al., 2016) and related article (Mansour L, Di Biase, Smith, Zalesky, & Seguin, 2023). The T1-weighted structural brain images were acquired using a 3D MPRAGE acquisition at 1 mm isotropic resolution with a 256 mm superior-inferior field of view.

Resting-state BOLD data were acquired with a multi-band gradient echo EPI sequence, with an acquisition time of 6 min, for a total of 490 volumes, with a spatial resolution of 2.4 mm isotropic voxels (TE/TR 39/735 ms, MB 8, no in-plane acceleration, flip angle 52°, conventional fat saturation)(Dollard & Miller, 1950) (Miller et al., 2016). Preprocessing steps (Alfaro-Almagro et al., 2018) consisted of the FSL MELODIC pipeline (Jenkinson, Beckmann Cf Fau - Behrens, Behrens Te Fau - Woolrich, Woolrich Mw Fau - Smith, & Smith) (EPI susceptibility distortion correction, gradient distortion correction, motion correction with FSL MCFLIRT, grand-mean intensity normalization, and high-pass temporal filtering) followed by an ICA + FIX step to suppress remaining artifact components. In addition to UKB quality control, participants with excessive head movement (average framewise displacement > 0.2mm, ID 25741) during scans were excluded.

Diffusion-weighted MRI (dMRI) data were acquired using a multi-band spin echo EPI sequence with a 7-minute acquisition time. The data were acquired across 100 unique diffusion sensitization directions, evenly distributed over two shells (b-values: 1000, 2000 s/mm^3^), along with 5 b=0 volumes. The spatial resolution was 2 mm isotropic voxels (MB=3, no in-plane acceleration, TE/TR=92/3600 ms, partial Fourier=6/8, conventional fat saturation). Additionally, 3 b=0 volumes were collected with reversed phase encoding to estimate susceptibility fields. Preprocessing including corrections for eddy currents, head motion, and gradient distortions (Alfaro-Almagro et al., 2018). Whole-brain tractography was estimated using *MRtrix3* (Tournier et al., 2019).

*Phenotype*

In our analysis we restricted usage to phenotype available at Instance 2 (i.e., the time-point where brain imaging data were collected). Items not available at Instance 2 (for example certain PHQ modules that were only administered at baseline) were therefore excluded from our analyses. Details of each ID and its questionnaire source are listed in the Supplementary Table 1.

*Cognitive function.* Cognitive items (Category 100026) included fluid intelligence score (ID 20016), maximum digits remembered correctly (ID 4282), prospective memory result (ID 20018), mean time to correctly identify matches (ID 20023), number of word pairs correctly associated (ID 20197), number of puzzles correct (ID 21004), and number of symbol digit matches made correctly (ID 23324).

*Mental health.* A wide range of items pertaining to mood, feelings, satisfaction, depression/anhedonia, mania/hyper, irritability, risk‐taking, and help‐seeking were included in Category 100060. Examples include: mood swings (ID1920), miserableness (1930), irritability (1940), sensitivity/hurt feelings (1950), fed‐up feelings (1960), frequency of depressed mood in last 2 weeks (ID 2050), ever depressed for a whole week (4598), ever manic/hyper for 2 days (4642), happiness (ID 4526; reverse‐scored in main text as "Unhappiness"), work/job satisfaction (4537), health satisfaction (4548), financial situation satisfaction (4581).

*Self-reported medical conditions.* We included variables (Category 1003). for example, the number of self‐reported cancers (ID 134) and number of self‐reported non-cancer illnesses (ID 135). We also included the number of treatments/medications taken (ID 137). Additionally, a range of condition-specific items were included: e.g., diabetes diagnosed by doctor, cancer diagnosed by doctor, age high blood pressure diagnosed, fracture/broken bones in last 5 years, hearing difficulty, chest pain, shortness of breath walking on level ground, etc. These variables were grouped in the main text under "Self-reported medical conditions". Other relevant self-report items included overall health rating (ID 2178; renamed in main text as "Overall unhealth rating" because higher values indicate worse health), long‐standing illness, disability or infirmity, falls in the last year, weight change compared with 1 year ago, various pain items (e.g., neck/shoulder pain for 3+ months; hip pain for 3+ months; back pain for 3+ months; knee pain for 3+ months; headaches for 3+ months) and sensory/vision (e.g., wears glasses/contact lenses, age started wearing them) and hearing (e.g., tinnitus).

*Lifestyle and Environmental Factors*

We extracted a broad set of self-reported measures from the UK Biobank baseline and online questionnaires to characterize multiple lifestyle and environmental domains. Most variables were collected through the UK Biobank baseline touchscreen questionnaires and online follow-up Mental Health Questionnaire (MHQ), launched in 2016. The childhood adversity questions (ID 20487-20491) originated from the Childhood Trauma Screener (CTS-5) module within the MHQ (see biobank.ndph.ox.ac.uk/ukb/label.cgi?id=145), while the broader adversity and trauma items were part of the Online Mental Health dataset (see ukbiobank.ac.uk). Dietary, lifestyle, and SES variables were derived from the baseline Touchscreen Questionnaire modules described on the UK Biobank data showcase. Details of each variable are provided in the Supplementary Table 1.

*Social support.* Measures included the number of people in household (ID 709; with "1" indicating living alone) and the frequency of friend/family visits (ID 1031).

*Socioeconomic status (SES).* SES was indexed by average total household income before tax (ID 738) and age completed full-time education (ID 845). In the main text, "Household" and "Education" were grouped under the SES category.

*Physical activity.* Variables included the number of days per week with ≥10 minutes of moderate (ID 884) or vigorous (ID 904) activity, average duration of moderate (ID 894) and vigorous (ID 914) activity, and time spent watching television (ID 1070) or using a computer (ID 1080).

*Sleep.* Sleep measures included sleep duration (ID 1160), with values < 6 hours classified as too short and ≥ 9 hours as too long (binary variables 1160_S and 1160_L), and sleeplessness/insomnia (ID 1200).

*Diet.* Dietary intake was assessed using frequency of consumption of major food categories, including cooked vegetables (1289), salad/raw vegetables (1299), fresh fruit (1309), dried fruit (1319), oily fish (1329), non-oily fish (1339), processed meat (1349), beef (1369), lamb/mutton (1379), pork (1389), bread (1438), and cereal (1458).

*Alcohol use and smoking.* Alcohol consumption was indexed by drinking frequency (ID 1558), weekly intake of specific beverage types (1568-1608), monthly intake counterparts (4407-4451), and drinker status (20117). Smoking status was assessed using ID 20116.

*Early life factor.* Early developmental measures included being breastfed as a baby (1677), adopted as a child (1767), multiple birth (1777), maternal smoking around birth (1787), and birth weight (20022).

*Trauma.* The trauma is screened using Childhood Trauma Screener (CTS-5, ID 20487-20531), including a series of items on childhood and adulthood adverse or supportive experiences, covering emotional neglect, physical abuse, and related domains. The broader set included questions on family relationships, partner violence, serious accidents, combat exposure, and other life-threatening or traumatic events.

*Preprocessing of genetic data*

Genotype calling was performed by Affymetrix (now part of ThermoFisher Scientific) on two closely related purpose-designed arrays. ~50,000 participants were run on the UK BiLEVE Axiom array (Resource 149600) and the remaining ~450,000 were run on the UK Biobank Axiom array (Resource 149601). The dataset combines results from both arrays (see Field 22000) and there are 805,426 markers in the released genotype data. The positions of markers in the data are in GRCh37 coordinates. It was not possible to assay genotypes for some participants (~3%) as sufficient DNA could not be extracted from their blood samples.

We restricted the sample set to participants who had undergone brain MRI scanning in our study, and analysed only autosomal variants (chromosomes 1-22). This approach ensures that the ancestry covariates included in the downstream GWAS reflect the actual sample analysed, and not the broader UKB sample whose ancestry structure may differ. Individuals with a genotype missingness rate >5% were excluded. Variants were filtered based on a minor allele frequency (MAF) threshold of 0.01, Hardy-Weinberg equilibrium (HWE) test p<1e-6, and variant missingness >5%. After QC, all 22 chromosomes were merged into a single data, allowing a maximum of two alleles per locus. To correct for population structure specific to the imaging subsample, we performed a principal component analysis (PCA) on the QCed merged dataset rather than using the population principal components provided by the UK Biobank.

*Genome-wide association studies (GWAS)*

GWAS analyses were conducted using PLINK 2.0, adopting an additive genetic model and a linear regression framework, in which allele dosage was modelled additively (0, 1, 2) for each SNP. The regression models included sex, age, BMI, head motion, SNR, TIV, centre, and the top 20 genetic PCs as covariates to account for potential confounding effects due to demographic or ancestry differences.

**Supplemental Figures**

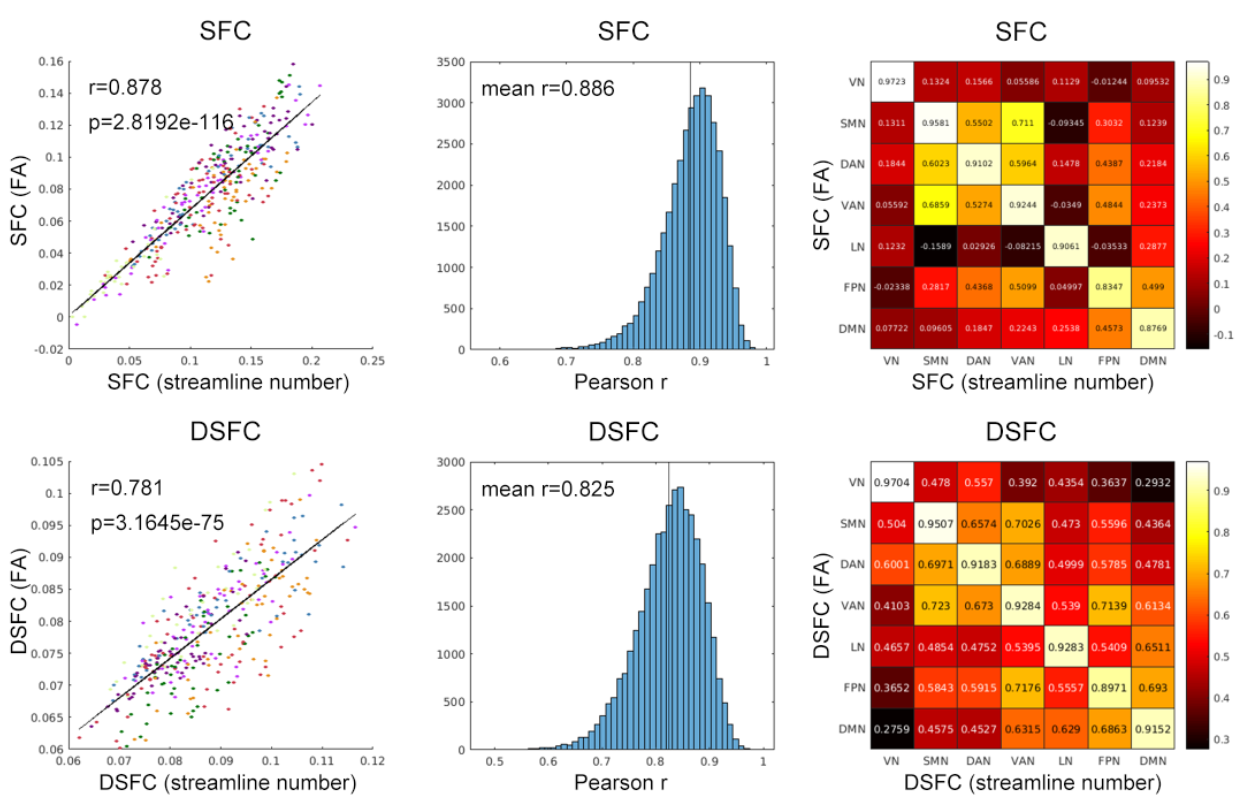

***Supplementary Fig. 1. Reproducibility of SFC and DSFC.*** *We repeated the calculation of SFC and DSFC using an alternative structural connectivity measure (the mean fractional anisotropy, FA, between two brain regions) with the same dynamic parameters, and found high reproducibility of our results across different structural connectivity measures. Left panel: Correlation between group-averaged streamline-based and FA-based SFC and DSFC brain maps (measured with Pearson's correlation coefficient, r). Color indicates functional networks (consistent with the Yeo7 network used in the main text). Middle panel: Histogram of the correlation (Pearson's r) between individual-level streamline-based and FA-based SFC and DSFC maps, assessing the spatial pattern reproducibility at the individual level. Right panel: Reproducibility of network-level streamline-based and FA-based SFC and DSFC brain maps, reflecting the similarity of SFC or DSFC across networks in the population, with Pearson's r displayed in the heatmap.*

**Supplemental Tables**

***Supplementary Table 1.*** *All variables used in this study, including the UK Biobank Category and Field ID, along with the Category labels used in the main text.*

***Supplementary Table 2.*** *Association between demographic variables (sex, age, body mass index/BMI, Intracranial Volume/ICV) and static structure-function coupling (SFC) and dynamic structure-function coupling (DSFC) (network-level mean). Reported statistics include the t-value (corresponding to the t-statistic of the β value), p-value, degrees of freedom (DF), beta (β-statistic), Cohen’s f², Cohen’s d, and confidence intervals (ci1: lower bound; ci2: upper bound), and similar for other tables.*

***Supplementary Table 3.*** *Association between demographic variables and SFC/DSFC (Glasser 360 parcellation).*

***Supplementary Table 4.*** *Association between cognitive, mental and physical health phenotypes and SFC/DSFC (network-level mean).*

***Supplementary Table 5.*** *Association between cognitive, mental and physical health phenotypes and SFC/DSFC (Glasser 360 parcellation).*

***Supplementary Table 6.*** *Association between environmental and lifestyle factors and SFC/DSFC (network-level mean).*

***Supplementary Table 7.*** *Association between environmental and lifestyle factors and SFC/DSFC (Glasser 360 parcellation).*

***Supplementary Table 8.*** *Mediation analysis results. Independent variable (X): Smoking Status. Mediator (M): SFC. Dependent variable (Y): Fluid Intelligence Score.*

***Supplementary Table 9.*** *Association between Polygenic risk score (PRS) and SFC/DSFC (network-level mean).*

***Supplementary Table 10.*** *Association between Polygenic risk score (PRS) and SFC/DSFC (Glasser 360 parcellation).*

***Supplementary Table 11.*** *AD-related genes from previous meta-analysis.*

***Supplementary Table 12.*** *AD-related SNPs significantly associated with SFC/DSFC (network-level mean).*

***Supplementary Table 13.*** *Association between APOE ε4 dosage and SFC/DSFC (network-level mean).*

***Supplementary Table 14.*** *Association between APOE ε4 dosage and SFC/DSFC (Glasser 360 parcellation).*

***Supplementary Table 15.*** *Mediation analysis results. Independent variable (X): Aging & APOE ε4 dosage. Mediator (M): SFC/DSFC. Dependent variable (Y): Fluid Intelligence Score.*

***Supplementary Table 16.*** *Validation analysis results. At network level, Pearson r between steamline-based SFC/DSFC and FA-based SFC/DSFC were calculated.*
